# Early vocal production and functional flexibility in wild infant chimpanzees

**DOI:** 10.1101/848770

**Authors:** Guillaume Dezecache, Klaus Zuberbühler, Marina Davila-Ross, Christoph D. Dahl

## Abstract

How did human language evolve from earlier forms of communication? One way to address this question is to compare prelinguistic human vocal behavior with nonhuman primate calls. Here, an important finding has been that, prior to speech, human infant vocal behavior exhibits functional flexibility, the capacity of producing protophones that are not tied to one specific function. Nonhuman primate vocal behavior, by contrast, is comparably inflexible, with different call types tied to specific functions. Our research challenges the generality of this claim, with new findings of flexible vocal behavior in infant chimpanzees. We used artificial intelligence consisting of automated feature extraction and supervised learning algorithms to analyze grunt and whimper vocalizations from free-ranging infants during their first year of life. We found that grunt production was highly flexible occurring in positive, neutral and negative circumstances, as already shown in human infants. We also found acoustic variants of grunts produced in different affective contexts, suggesting gradation within this vocal category. By contrast, the second most common call type of infant chimpanzees, the whimpers, was produced in only one affective context in line with standard models of nonhuman primate vocal behavior. We concluded that the most common chimpanzee vocalization, the grunt, qualifies as functionally flexible, suggesting that evolution of vocal functional flexibility occurred before the split between the Homo and Pan lineages.

## INTRODUCTION

At some point in evolutionary history, there must have been a transition from primate-like inflexible to human-like flexible acoustic communication, which may have coincided with the origins of speech. The evolutionary history of this transition continues to be vividly debated (Fitch, 2018), with a large range of comparative evidence from animal communication systems, with the consensus view that direct evolutionary homologies are generally absent in the primate order (Rendall & Owren, 2002). More recently, however, some vocal and neural equipment has been identified in different primate species that allow for the production of speech-like sounds (Boë et al., 2017; Fitch, Boer, Mathur, & Ghazanfar, 2016; Lieberman, 2017) and for limited control over vocal fold oscillation (Lameira & Shumaker, 2019).

A similar point of contention is whether there are fundamental differences in the ontogenetic trajectories between non-human primate and human vocal behavior prior to speech. By the age of one-month, human and great ape infants share parts of their vocal repertoire, such as crying or laughter, suggesting a common evolutionary root and biological function insofar as, in both species, the calls possess an illocutionary quality that expresses intuitively identifiable internal states to caregivers and other listeners (Jhang & Oller, 2017; Oller et al., 2013).

However, human infants also produce a range of other sounds (called ‘protophones’), such as squeals, vocants and growls, that are not tied to particular affective states (Jhang & Oller, 2017; Oller et al., 2013). Some of these pre-linguistic sounds may convey relatively specific meanings, in the sense that infants produce them to express specific behavioral intentions rather than communicating specific affective states (Kersken, Zuberbühler, & Gomez, 2017). This apparent decoupling of signal structure and function in young infants, termed ‘vocal functional flexibility’, has been identified as a major evolutionarily precursor to language (Oller et al., 2013). Because of its early ontogenetic onset, vocal functional flexibility is said to be more foundational to human speech than other building blocks of the language faculty, such as proto-syntax or vocal elaboration (Oller et al., 2013). Vocal functional flexibility, in this view, is a prerequisite for speech development, and a major evolutionary departure from the functionally inflexible vocal behavior of non-human primates (Waal & Pollick, 2011).

In one relevant study, Clay et al. (Clay, Archbold, & Zuberbühler, 2015) examined ‘peep’ calls in mature bonobos (*Pan paniscus*), their most common vocalizations, and found that they are produced in a variety of contexts, ranging from seemingly positive (food provisioning) to neutral (travel and resting) to negative (agonistic and alarm) situations. Based on these findings, the authors concluded that bonobos have the capability to produce sounds that are not affectively biased (Clay et al., 2015). Their peeps were, however, attributed to broad behavioral contexts (such as feeding or travelling) with no focus on more specific and transient behaviors that may infer affective contexts, such as when individuals suddenly experience aggression during travelling and feeding bouts. As such, the bonobo data are indicative of their peeps occurring across broad behavioral contexts but ultimately remain inconclusive in regards to whether vocal functional flexibility is indeed present in species other than humans. A second study, also on bonobos (Oller et al., 2019), suggests protophone-like vocal behavior with infants producing calls that occur in both low or moderate arousal situations, implying no affective binding. This conclusion has been preliminary, however, for the affective quality of the contexts surrounding vocalizations has proven difficult to discern.

Here, we directly addressed the functional flexibility hypothesis by examining infant chimpanzee (*Pan troglodytes schweinfurthii*) vocal behavior at a very early age (< 12 months) and their affective context of occurrence. Examination of early vocal production is critical for a more direct comparison with findings on human infants (Oller et al., 2013) and to test hypotheses about the evolutionary origins of functionally flexible vocal behavior. We took advantage of recent developments of applying machine learning techniques to the study of animal communication. We focused on two call types, the grunts and the whimpers, as they are acoustically distinct vocalizations that are common in young infants. Grunt calls are of particular importance as they develop into a central component of the vocal repertoire of chimpanzees and contribute to a variety of vocal sequences produced by juveniles, sub-adults and adults (Crockford & Boesch, 2005). For example, grunts complement panting elements during laughter and when encountering dominant individuals (‘pant-grunts’). They are also produced upon encountering a food patch or when joining a foraging party (‘rough grunts’). Finally, they are routinely produced throughout resting or in relaxed social activities (Goodall, 1986).

Like in humans (McCune, Vihman, Roug-Hellichius, Delery, & Gogate, 1996), grunts are produced from the first days of life in chimpanzees. Their ontogenetic development has already been studied to some degree in chimpanzees, which has shown some flexibility in usage (Laporte & Zuberbühler, 2011). Two types of grunts can be distinguished, although no study has yet offered an acoustical validation of the existence of these diverse types. First, uh-grunts are short, tonal sounds, resembling human vowels {u}, {o} and {a}, sometimes produced in short series (staccato-grunts) (Kojima, 2003; Plooij, 1984). The second type are the so-called ‘effort’ grunts, which represent the majority of grunting behavior in immature chimpanzees and are also present in humans and other mammals (McCune et al., 1996). Effort grunts are relatively soft and noisy and occur mostly during locomotor activities (Plooij, 1984). However, in adults, they are not mere byproducts of locomotion since they are sometimes emitted in the absence of movements, suggesting that the calls go through an ontogenetic transition from mere by-products of mechanical efforts to functionally active communicative signals in adults.

Another common vocal utterance produced by chimpanzee infants is whimpers (Dezecache, Zuberbühler, Davila-Ross, & Dahl, 2019; Levréro & Mathevon, 2013; Plooij, 1984). They are short, tonal and often produced in series with an upward shift in fundamental frequency. Contrarily to grunts, whimpers preferentially occur in aversive contexts, likely homologous to human crying or distress calls in other mammals (Plooij, 1984). Previous research (e.g., (Plooij, 1984)) has suggested the presence of whimper subtypes (single, serial and human-like whimpers), but we are not aware of any systematic acoustical analysis that would justify this nomenclature.

To address the hypothesis that vocal flexibility in grunts evolved before the split between *Pan* and *Homo* lineages, we examined the vocal behavior of six wild chimpanzee infants aged between 0-12 months old from the Sonso community of Budongo Forest, Uganda. We analyzed the extent to which vocal production of grunt-like and whimper-like vocalizations were affectively biased, i.e., occurring in positive, negative or neutral situations.

## RESULTS

### Types of vocal utterances

We inspected N = 1,016 vocal occurrences, of which N = 967 could be classified as either ‘grunts’ (N = 833) (corresponding to a rough, harsh and noisy sound) or ‘whimpers’ (N = 134) (usually a series of low-pitch tonal calls with increase in fundamental frequency throughout the series). Other types of calls were identified as ‘hoos’ (n = 23), ‘pants’ (n = 15), ‘screams’ (n = 2), ‘squeaks’ (n = 2) or ‘barks’ (n = 4). ‘Laughter’ (defined as grunting and panting) was uncommon (n = 3).

### Functional flexibility

Grunts: 44.8% of grunt-like vocalizations co-occurred with contexts classified as ‘positive’, 40.9% with ‘neutral’, and 14.3% with ‘negative’. When considering each individual separately, a similar picture emerged (Figure 1), with most grunt-like vocalizations co-occurring with ‘positive’ and ‘neutral’ affective contexts. We sought to evaluate the evenness of the distribution of grunts across affective contexts and did so by calculating, for each infant, the numerical dominance of one affective context over the others (from 1 / 3 [= equiprobability of all 3 affective contexts] to 1 [= complete dominance of one of the affective contexts over the two others]). We found dominance to be relatively low in grunts, varying from 0.37 and 0.63 (mean = 0.53; SD = 0.10), suggesting a stable and relative evenness in the affective distribution of grunts.

**Figure 1.**
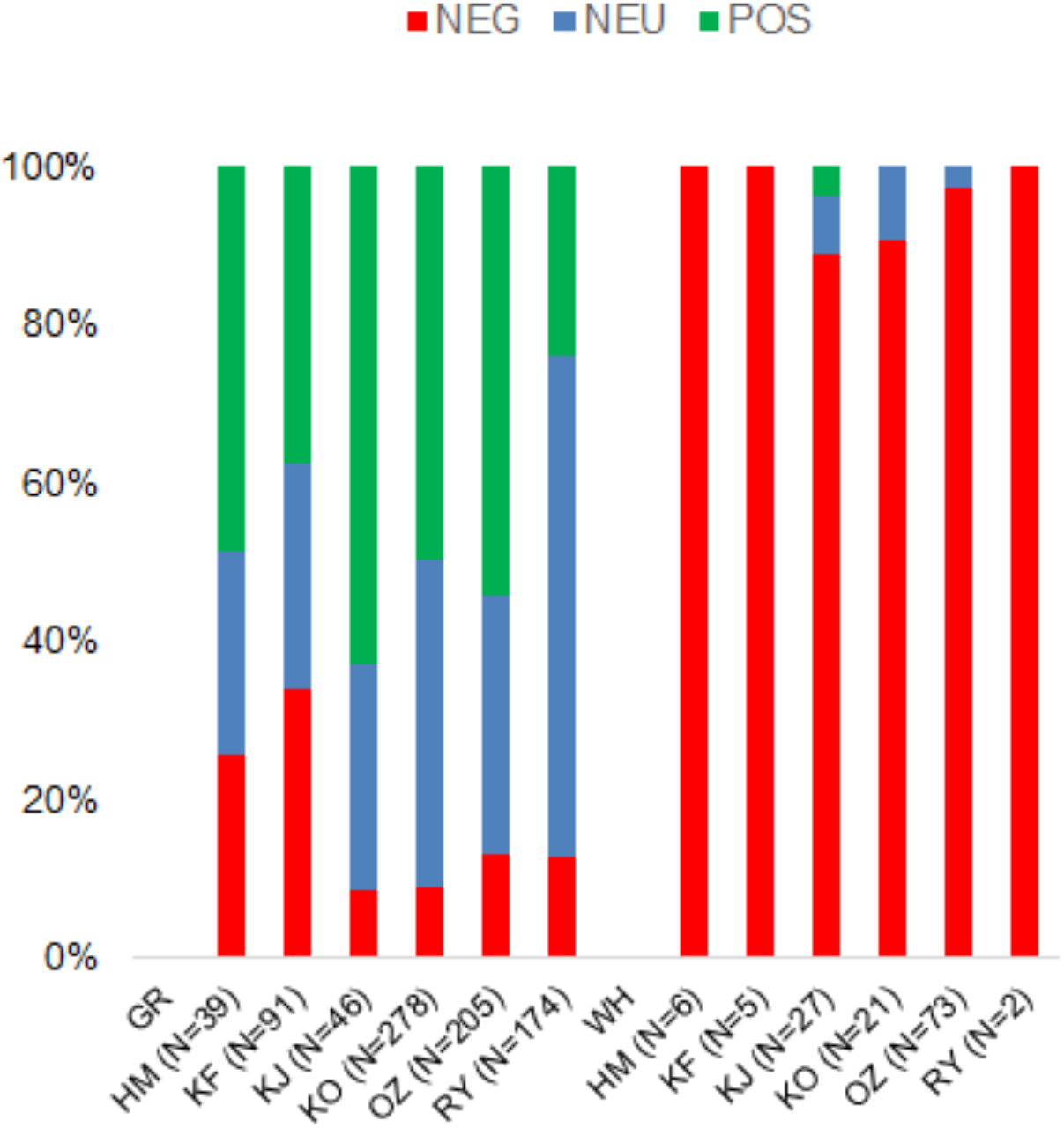
Proportion of grunt-like (GR) and whimper-like (WH) vocal behaviors recorded in negative (NEG), neutral (NEU) and positive (POS) affective contexts, for each individual separately. Number between brackets indicate the number of GR and WH calls contributed by each individual.

Whimpers: 94.8% of whimpers co-occurred with negatively classified contexts, and rarely with neutral (4.5%) or positive (0.7%) affective contexts. Inspection of individual distributions revealed the same pattern with whimper-like vocalizations systematically co-occurring with negatively classified contexts (Figure 1). The dominance of one affective context over the others in whimpers was relatively high, ranging from 0.89 to 1 (mean = 0.96; SD = 0.05), indicating low evenness in the affective distribution of whimpers.

Grunts vs. Whimpers: When comparing the distributional evenness of grunts vs. whimpers, we found dominance to be statistically higher in whimpers than in grunts (paired Wilcoxon signed rank test: V = 21, *p* = .031).

### Acoustic variants of grunts

We then classified N=180 grunts (N=60 per affective context) according to their association with positive, neutral, negative contexts in order to test for the presence of acoustic variants. In the first step, we followed a feature extraction procedure by extracting the means and covariances of mel frequency cepstral coefficients (MFCCs) for each call, and compared these values according to the calls’ associations (e.g. positive vs negative) using *t*-tests. This approach provides a general idea of how well positive, neutral and negative calls can be separated. We displayed the resulting *p*-values in an empirical cumulative distribution function (eCDF) (Figure 2A). We found that 5-10% of all features showed significant differences between the class labels at a 5%-significance level. In other words, 5-10 of 104 feature dimensions had strong discrimination power to distinguish between grunts pertaining to the various affective contexts.

**Figure 2.**
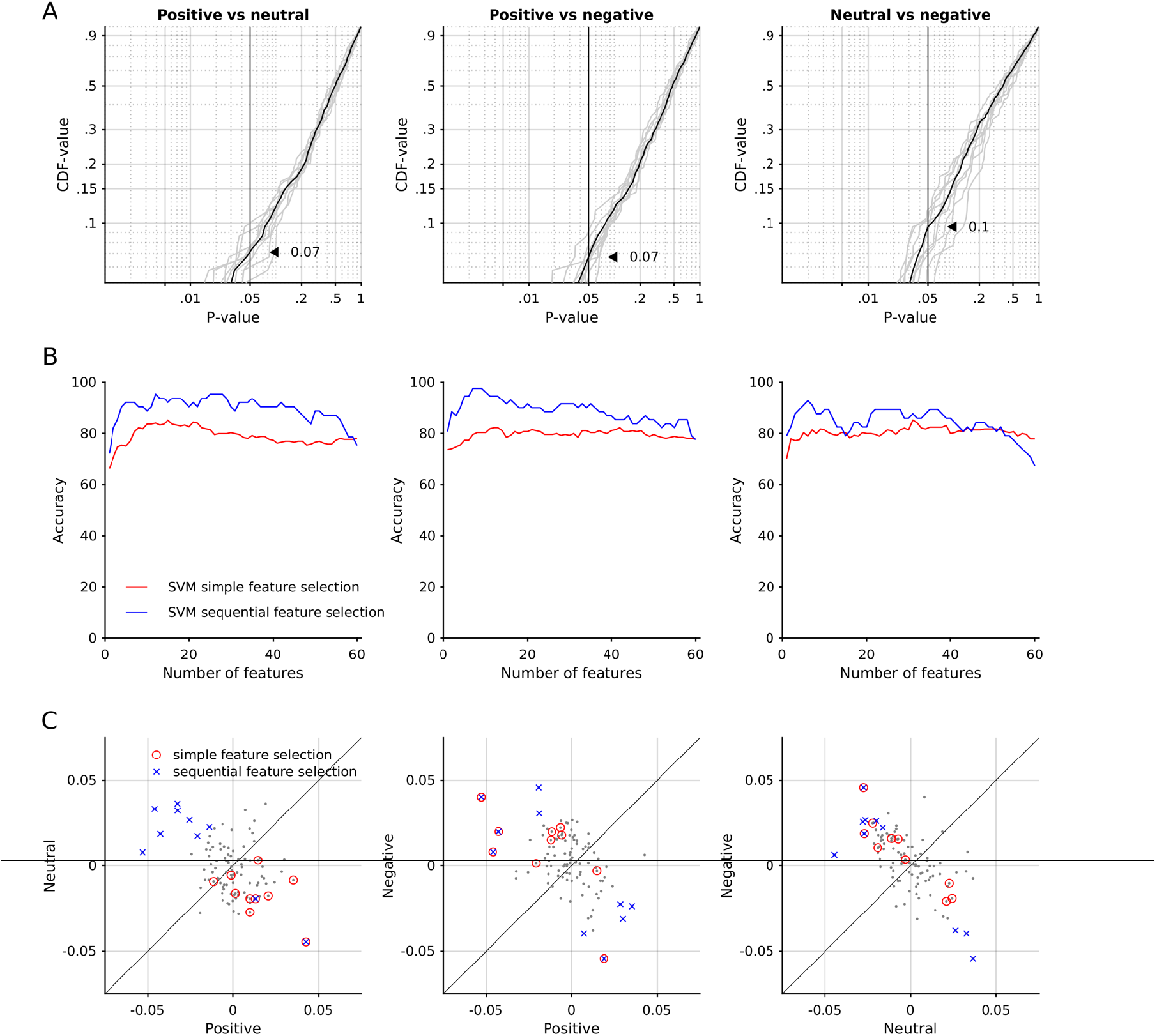
Feature selection and classification performances. The columns represent the comparisons of affective contexts during which the vocal utterance occurred. A. For each feature dimensions the discrimination power of the two classes (e.g. positive vs. neutral) was evaluated using a t-test. P-values are shown as an empirical cumulative distribution function (eCDF). Gray lines show the results of individual runs of evaluation; black lines show the means of individual runs. Indicated with arrow heads are the proportions of feature dimensions that significantly discriminate between the two classes tested. B. The classification performances are shown for the SVM classifier relying on feature dimensions extracted through a simple feature selection (red lines) and a sequential feature selection procedure (blue lines). C. Feature selection outcomes are shown for simple (red circles) and sequential feature selection (blue x-s) as overlays on all feature dimensions (black dots).

In the second step, the feature selection procedure, we systematically varied the number of feature dimensions to be considered into the classification process (x-axis in Figure 2B). The feature dimensions that went into the classification process were determined by means of two methods: (1) simple filter feature selection method and (2) sequential feature selection method.

With the simple feature selection algorithm, the SVM correctly discriminated between classes at up to 80% (positive vs neutral: M = 78.99, SD = 3.53, *t*(59) = 63.69, *p* < .001; positive vs negative: M = 79.58, SD =1.83, *t*(59) = 125.37, *p* < .001; neutral vs negative: M = 80.44, SD = 2.06, *t*(59) = 114.26, *p* < .001; red lines in Figure 2B). A substantial improvement was found when sequentially selecting feature dimensions: SVM correctly classified samples at up to 95% (positive vs neutral: M = 89.56, SD = 4.84, *t*(59) = 143.42, *p* < .001; positive vs negative: M = 88.72, SD = 4.49, *t*(59) = 153.11,*p* < .001; neutral vs negative: M = 84.27, SD = 5.23, *t*(59) = 124.91, *p* < .001; blue lines in Figure 2B). For all comparisons chance levels were 50% due to the two-class comparisons applied. We further illustrated the simple feature selection outcomes by highlighting the feature dimensions selected (red circles in Figure 2C) among the feature dimensions not selected (black dots). Further, the features selected via the sequential feature selection are marked with blue x’s. It becomes evident that the sequential feature selection yields better performance through sequential combinations of feature dimensions that, on average, fall more distal to the diagonal midline than the feature dimensions selected by the simple feature selection process. Sequential feature selection, to a large extent, included feature dimensions not selected by the simple feature selection method.

We further ensured that each individual was not contributing solely to the classification results of various contrasts. To test this, we repeated the classification process for the number of individuals (N = 6) and excluded one individual at each time. As can be seen in Supplementary Figure 1, the classification performance did not improve nor deteriorate systematically, suggesting no effect due to caller identity (the average *t*-value of one-sample *t*-tests is 97.52 +/− 30.25 (SD); all *p*-values were smaller than .001).

The use of means and covariances of cepstra yielded relatively high-performance scores in the classification routines at low computational loads. To assess whether certain feature dimensions (means and covariances of cepstra) occurred above chance across all comparisons, we determined the empirical distribution of occurrences of feature dimensions and contrasted it with a random distribution. While the use of the same feature dimension in up to 33% of the comparisons was not significantly different in the empirical distribution from the random distribution, the use of the same feature dimension in 50% of comparisons was significantly increased in the empirical distribution (Figure 3A). To describe the frequency bands explaining significant variances between classes of calls, we traced back the frequency bands underlying the significant feature dimensions, i.e., covariances of cepstra, and determined the sign of the covariances. We found a negative covariances between the following frequency bands (Figure 3B): (1) band 2 (196.30 to 488.89 Hz) and band 4 (488.89 to 927.78 Hz), (2) band 4 (488.89 to 927.78 Hz) and band 8 (1074.07 to 1366.67 Hz), band 6 (781.48 to 1074.07 Hz) and band 9 (1220.37 to 1512.96 Hz). We found a positive covariance between the frequency bands 9 (1220.37 to 1512.96 Hz) and 10 (1366.67 to 1659.26 Hz). Mean cepstra were significantly contributing in the frequency bands from (1) 50 to 342.59 Hz, (2) 196.30 to 488.89 Hz, (3) 927.78 to 1220.37 Hz.

**Figure 3.**
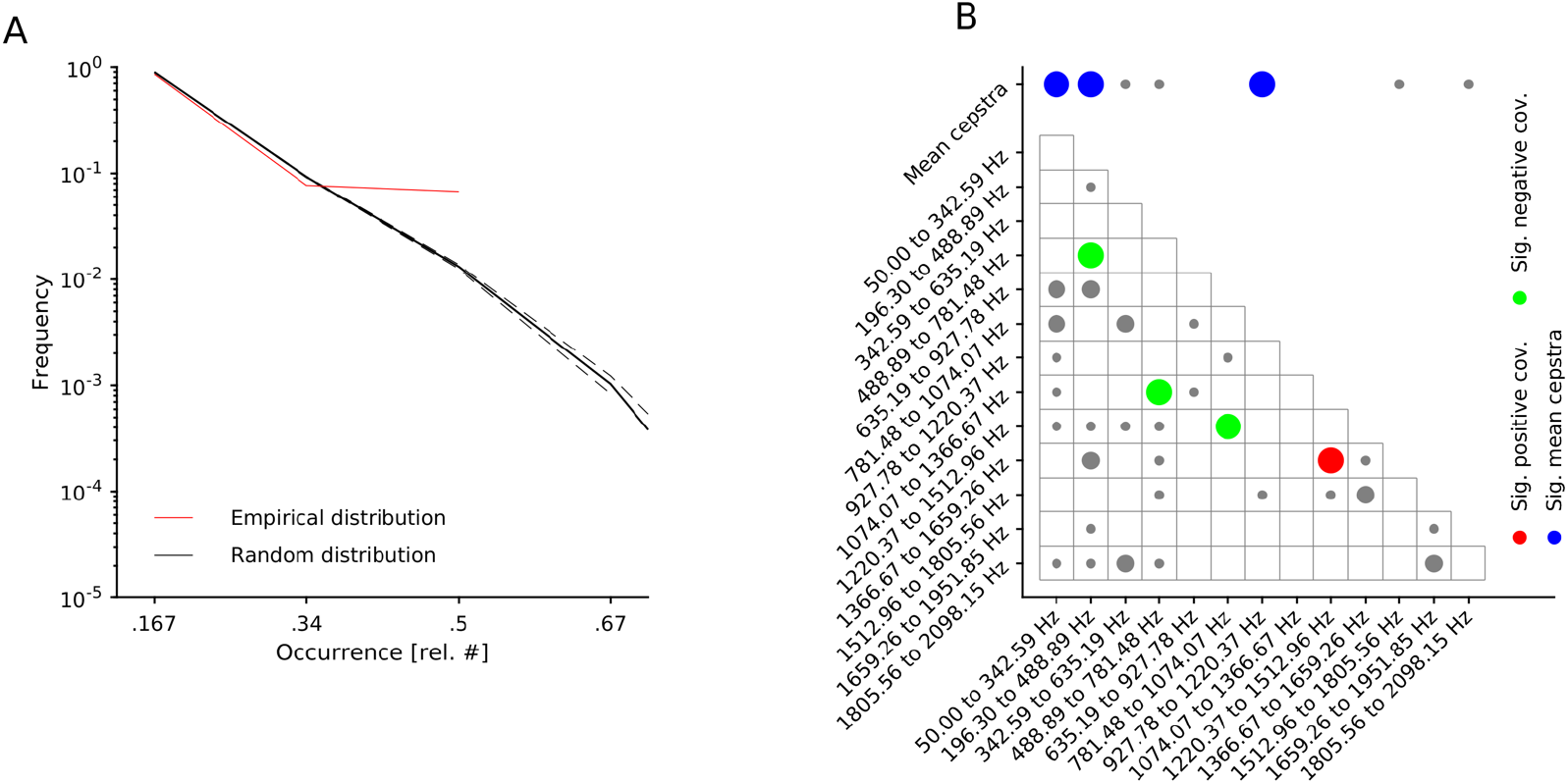
Overall feature importance. A. The empirical distribution of feature dimensions across all comparisons. B. Significant feature dimensions are shown in colors, according to their sign: in red positive covariances, in green negative covariance. The means of cepstra are shown in blue. The marker size indicates the occurrence: small = 1, medium-large = 2, large = 3 (significant). Gray-colored markers are non-significant feature dimensions.

## DISCUSSION

Oller and colleagues (Jhang & Oller, 2017; Oller et al., 2013; Oller, Griebel, & Warlaumont, 2016; Oller & Griebel, 2004) posit that speech emerged from pre-linguistic vocalizations that are free of predetermined biological function, a precursor called ‘vocal functional flexibility’. Modern human infants regularly vocalize in such a way, in supposed contrast to the relative inflexibility of vocalizations in non-human primates (e.g., (Pollick & Waal, 2007; Waal & Pollick, 2011)). Functionally flexible vocalizations of young human infants, according to this theory, have evolved in humans in relation to allo-maternity (Burkart, Hrdy, & Van Schaik, 2009; Hrdy, 2007, 2009; Kramer, 2010; Schaik & Burkart, 2010) or altriciality (Locke, 2006) and associated pressures on young infants to signal their needs and attract caregivers (Ghazanfar, Liao, & Takahashi, 2019; Locke, 2006; Zuberbühler, 2012).

In the current study, we focused on the grunt-like and whimper-like calls of young chimpanzee infants, using novel coding strategies and state-of-the-art acoustic analysis tools. By contrast to previous studies, we elaborated a workable coding system which provides insight into the affective state of the animal, without solely relying on the broad behavioral contexts. We found that grunt-like calls are functionally flexible vocal units, produced frequently by chimpanzee infants in both positive and neutral situations, and less commonly also in negative situations. Importantly, the presence of grunts in contexts of low-to-mild arousal is consistent with the hypothesis of vocal functional flexibility (Oller et al., 2019), and so is the finding that grunts occur in similar proportion in positive and neutral contexts (Oller et al., 2013). On the other hand, whimper-like vocalizations seem to be confined to situations involving negative affective states in the infants. Their near absence in positive and neutral situations suggests that they represent a functionally rigid vocalization that has evolved for a narrow range of biological purposes, similar to cries in humans (Oller et al., 2013), to which they may functionally correspond (Goodall, 1986). Grunts, more generally, are a promising class of calls, insofar as their functional flexibility is in line with the ubiquity of this vocal category in a diversity of contexts in other primate species (Cheney & Seyfarth, 1982, 2018; Range & Fischer, 2004; Rendall, Seyfarth, Cheney, & Owren, 1999; Salmi, Hammerschmidt, & Doran-Sheehy, 2013). This being said, data suggest that production of sounds that are typically uttered under high stress (e.g., alarm calls) can also occur in the absence of the triggering stimulus (Lameira & Call, 2018), a pattern also suggested by the use of alarm calls to deceive conspecifics during foraging activities (Møller, 1988; Wheeler, 2009).

Our second finding was systematic acoustic differences between grunts given in positive, neutral and negative situations, which enabled us to segregate acoustic variants of grunts into these categories. Acoustical differences linked to the affective context surrounding vocal production are common in humans as in other animals (Arias, Belin, & Aucouturier, 2018; Aucouturier et al., 2016; Banse & Scherer, 1996; Briefer, 2012; Goupil, Johansson, Hall, & Aucouturier, 2019; Ponsot, Burred, Belin, & Aucouturier, 2018; Williams & Stevens, 1972). Our data suggest that there is inter-gradation between grunt-types, with differences in acoustics relating to differences in contexts. Grunts, in other words, represent a coherent and unified call type that can manifest itself in acoustic variants in relation to the affective contexts in which they are produced.

How exactly functionally flexible vocalization produced by human infants transition into speech sounds has been described in previous studies (Boysson-Bardies, 2001; de Boysson-Bardies, 1993; de Boysson-Bardies & Vihman, 1991; Elbers & Ton, 1985; Nathani, Ertmer, & Stark, 2006; Oller, 2000; Oller, Wieman, Doyle, & Ross, 1976). Although chimpanzee infants produce grunts in ways consistent with the functional flexibility hypothesis, they of course never produce speech sounds and, historically, have failed to acquire human speech utterance even after extensive training (Hayes & Hayes, 1951). Instead, infant chimpanzee grunts may gradually develop into context-specific call variants with seemingly relatively narrow biological functions (Laporte & Zuberbühler, 2011; Slocombe & Zuberbühler, 2010; Slocombe & Zuberbühler, 2005; Watson et al., 2015), with clear acoustical boundaries notably between grunts used in greeting situations (‘pant-grunts’) and those produced upon encountering food. It is possible that the acoustic boundaries we identified between the grunts produced across affective states are the foundation of acoustic diversification in adults, although the categories used to define the affective states here (for instance, feeding and social approach are together considered ‘positive’) are not consistent with the vocal differentiation seen in adults (the grunts produced in feeding vs. social approach situations are acoustically distinct in adults (Crockford, in press; Goodall, 1986)). Alternatively, those calls may simply disappear and be absent from the adult repertoire, one causal factor being the relative absence of social reinforcement (including contingent vocal responses (Ghazanfar et al., 2019)) associated with grunt production, as compared to the frequent maternal reactions to distress calls (Dezecache et al., 2019).

Our tentative to further explore the affective state of the infant by considering other cues, such as the infants’ facial expressions or the mothers’ behavior, faced considerable challenges. We found that infant facial movements are extremely fast and fluid, which prevented us from reliable coding particularly in the wild. For this reason, the behavioral context of the infant alone was the most relevant available cue to approach the affective dimension of the situation. While we must yet acknowledge the limitations pertaining to the fact that judgments of infants’ affect were made based on the infants’ behavioral contexts and done so by a human observer, the results of the acoustic analysis are providing important support for the approach used to categorize affect in the present work. Future studies should investigate the affective impact of other communicative signals used by infants (Fröhlich & Hobaiter, 2018; Fröhlich, Wittig, & Pika, 2018).

Another hint to the affective dimension of the situation is the mothers’ behavior. Protocols where mothers may be asked to interact with toddlers may yield to responsiveness from the mothers whichever the affective state of the infant is (Yoo, Bowman, & Oller, 2018). In the course of spontaneous behavior, though, we may expect little intervention from the chimpanzee mothers, except in situations where the infant is in danger. In our sample, responsiveness of the mother (tentatively defined in pilot coding as being either proactive, protective or neutral by the observer) was relatively low (proactive or protective less than half of the time), a pattern which might be due to differences in mothering style between chimpanzees and humans, or a difference between our own study (where no particular demand is put on the mother) and others (where mothers may be interacting with their infant, e.g., (Oller et al., 2013)).

Although playback of infant grunts to the mother may appear like a methodological possibility to further establish maternal assessment of the affective state of the infant or to be able to see whether mothers respond differently to positive, neutral and negative grunts (Fischer, Noser, & Hammerschmidt, 2013; Fischer, 2016; Zuberbühler, 2014), this would require either playing the infants’ calls in its own presence (which is ethically inappropriate) or playing the calls of another infant to a mother (which may not trigger any reaction at all in the non-genetically related mother).

In latest research, the comparative volubility (quantity of sounds produced in a given period of time) of human infants and other animals (Ghazanfar et al., 2019; Ghazanfar & Takahashi, 2014; Oller et al., 2019; Takahashi et al., 2015), and the privileged function of protophone-like vocalizations to elicit social interactions and vocal turn-taking with caregivers (Oller et al., 2019; Yoo et al., 2018). In humans, non-affectively bound vocalizations appear to occur more often than affectively bound vocalizations (such as crying) (Oller et al., 2019). They occur in solitary contexts where infants invest in practice and vocal exploration. They also occur in interactive contexts, so as to elicit and regulate social interactions with caregivers. Caregivers appear to detect the functional difference between protophones (as potentially interactive calls) and other calls (such as cries), where caregiver intervention is solicited (Yoo et al., 2018). Comparison with bonobo infants suggested much higher rate of production of non-affectively bound vocalizations and much higher vocal investment in social interactions in human infants (Oller et al., 2019). Whether human infants also are comparably more talkative than their chimpanzee counterparts is a question we need to be exploring. This should be preferably investigated in captive or semi-captive settings, where true calling rate can be assessed, for video monitoring is less likely to be interrupted and for levels of ambient noise could be comparatively less problematic. Such problems have already been acknowledged by (Oller et al., 2019) regarding previous report on the flexible development of grunting behavior in wild chimpanzees as well as their rate of occurrence (Laporte & Zuberbühler, 2011). Data from the vocal development of one captive chimpanzee indicate lower volubility than in humans (Kojima, 2003). Future studies should evaluate this fact with a larger sample.

In conclusion, our study suggests that chimpanzees may possess a feature that is fundamental to the development of speech in humans, the ability to produce vocalizations that are not strongly bound to one particular affective context, but are produced in a functionally flexible manner. However, we should expect that future research will reveal further examples. For instance, coo calls in several macaque species (Hsu, Chen, & Agoramoorthy, 2005; Owren & Casale, 1994) or grunts in vervet monkeys (Seyfarth & Cheney, 1984) also seem to be given in a variety of contexts. Future research will equally have to address the question of how selection favored acoustic diversification of functionally flexible vocal behavior into speech in humans. The main driver for this transition, it has been argued, may have been the highly cooperative breeding system of humans, with infants regularly looked after by individuals other than the mother, which requires infants to become more active agents in forming social bonds from a much younger age than in great ape infants (Ghazanfar et al., 2019; Zuberbühler, 2012).

Cooperative breeding, in this view, may thus have transformed a functionally flexible vocal system, evolved prior to the split between humans and apes, into the uniquely human way of using vocal signals to interact socially. Another complementary reasoning is that humans’ high altriciality selected for the most vocal individuals, capable of attracting caregivers (Locke, 2006). Future studies should clarify the relative contribution of both factors through mapping the phylogenetic distribution of vocal functional flexibility.

## METHODS

### Ethics

Permission to conduct the study was obtained from the Ugandan Wildlife Authority (UWA) and the Uganda National Council for Science and Technology (UNCST).

### Subjects and data collection

Data were collected in the Sonso community of the Budongo Forest Reserve, Uganda (Reynolds, 2005) between February-June 2014, December 2014 and March-June 2015. This community comprises around 70 individuals well habituated to human observers. The natural behavior of N=7 infants was video recorded continuously during focal animal sampling, using Panasonic HC X909/V700 cameras, with a Sennheiser MKE-400 shotgun microphone. Six of those infants produced enough calls to be further considered in the statistical analysis (see Table 1 for details).

**Table 1.** List of focal animals, with their name (ID), sex and minimum and maximum age in months. Also given are the number of grunt-like and whimper-like vocal behaviors collected, as well as grunt-like vocalizations acoustically analyzed.

| *ID* | *Sex* | *Min. Age*<br>*(in*<br>*months)* | *Max. Age*<br>*(in*<br>*months)* | *N whimper-like*<br>*vocalizations* | *N grunt-like*<br>*vocalizations* | *N of grunt-like*<br>*vocalizations used in*<br>*acoustical analysis* |
| --- | --- | --- | --- | --- | --- | --- |
| HM | F | 3.41 | 6.85 | 6 | 39 | 10 |
| KF | M | <1 | 11.87 | 5 | 91 | 20 |
| KJ | M | 6.98 | 10.52 | 27 | 46 | 7 |
| KO | M | 3.08 | 8.46 | 21 | 278 | 67 |
| OZ | M | 1.38 | 8.16 | 73 | 205 | 32 |
| RY | M | 4.75 | 8.16 | 2 | 174 | 44 |

### Behavioral data analysis

Videos were inspected for the presence of infant vocalizations. We defined vocal behavior as the occurrence of single sound units or series of sounds produced by the infant’s vocal apparatus, separated by a least 5 seconds of silence.

As of today, there is no definitive repertoire of infant chimpanzee vocal behaviors. The categories used in this research are based on GD’s assessment. This assessment proved reliable when confronted to an independent assessment with Derry Taylor, using vocalizations from infant and juvenile semi-wild chimpanzees from the Chimfunshi Wildlife Orphanage, Zambia, collected by DT. One hundred-and-sixty vocalizations were indeed classified as belonging to either the ‘grunt’, ‘whimper’, ‘scream’ or ‘laughter’ category. Agreement was excellent (k = 0.77), even better when considering only ‘grunts’ and ‘whimpers’ (k = 0.92).

For each vocal occurrence, we coded infant behavior from the following list of mutually exclusive behavioral contexts (summarized in Table 2). The internal state of the infant was classified as ‘positive’ if it showed one of the following four behaviors: (1) ‘play’ (showing relaxed, joyous movements without obvious purpose, either as ‘social play’(accompanied by tell-tale behavior such as embracing and gentle biting) or ‘solitary play’; (2) giving or receiving ‘grooming’ (note that allo-grooming was never observed in our infants); (3) ‘feeding’ (breastfeeding or swallowing an edible element), and (4) ‘social approach’ (greeting a conspecific while moving towards it).

**Table 2.**
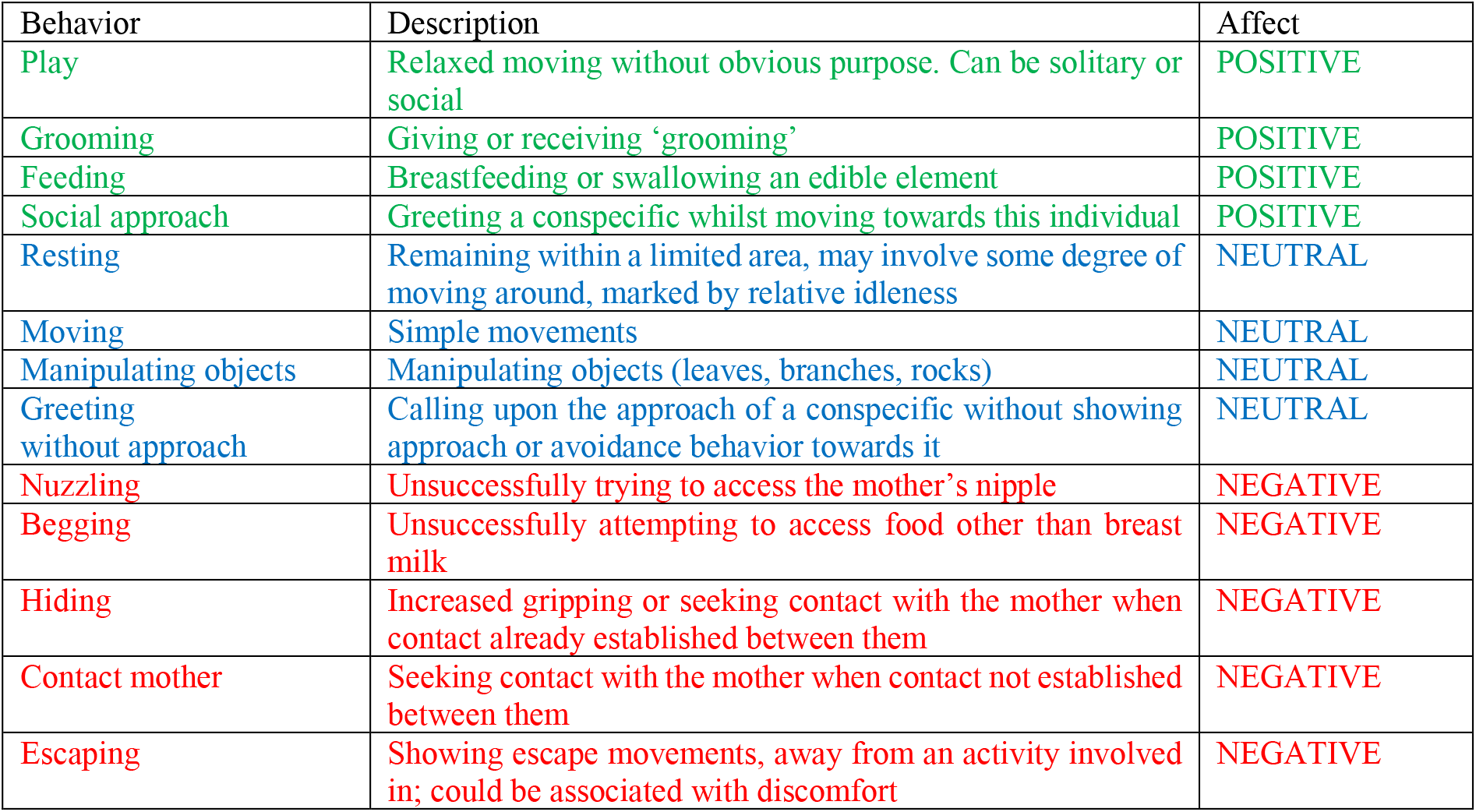
Affective coding of infant behavior

The infant’s internal state was classified as ‘neutral’ if it showed one of the following behaviors: (5) ‘resting’ (remaining with a limited area, sometimes moving within); (6) ‘moving’; (7) ‘manipulating objects’ (such as leaves, branches, rocks) without playful postures, or (8) ‘greeting without approach’ (calling upon the approach of a conspecific without showing nor approach or avoidance behavior towards it).

Infant behavior was classified as ‘negative’ if it showed one of the following behaviors: (9) ‘nuzzling’ (unsuccessfully trying to access the mother’s nipple); (10) ‘begging’ (attempting to access food other than breast milk); (11) ‘hiding’ (increased gripping or seek for contact with the mother when contact was already established between them); (12) ‘contact mother/kin’ was coded if infants were urgently seeking contact with the mother or a kin when contact was not already established between them; (13) ‘escaping’ (when the infant shows escape movements away from an activity it is involved in). Escaping could also include moment of discomfort when the infant is pressed against the belly of the mother or stuck in a bad position.

We performed intra-coder reliability tests on the affective contexts coded as positive, neutral and negative. For this, we randomly selected 200 video clips (around 19% of the coded dataset composed of the 7 infants), which were coded independently during two coding sessions more than a year apart (November 2015 and February 2017), so that the second coding was, notably, naïve. We found strong agreement between the two coding sessions (k = 0.73).

### Statistical analyses

In order to evaluate the evenness of the distributions of grunts and whimpers across affective contexts, we calculated, for each infant, and for grunts and whimpers separately, the dominance of one affective context over the two others, using the Berger-Parker Dominance index:

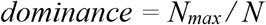

where N_max_ is the number of calls in the most abundant affective context; N the total number of calls across all affective contexts. Dominance values range from 1 / number of affective contexts (= equiprobability of calls across affective contexts; here 1 / 3 = 0.33) to 1 (= complete dominance of one of the affective contexts over the others).

Dominance values were compared between grunts and whimpers using a paired Wilcoxon Sign-Ranked test. Analyses were carried out using R (version 3.6.1 (R Core Team, 2018)) and R Studio (version 1.2.1335 (RStudio Team, 2015)).

### Acoustic analyses

Acoustic data analysis focused on grunts for they were the only vocal category for which at least two of the affective contexts were well represented. The acoustic structure of whimpers has been analyzed as part of another study. N=180 grunts were extracted. For each affective context, 60 were randomly selected. Following extraction, we used MATLAB (MathWorks Inc., Natick, MA, USA) for the acoustic data analysis, consisting of features extraction, feature selection and call classification. We first pre-processed the audio files by applying a band pass filter from 50 to 4000 Hz and normalized the signals using the following function:

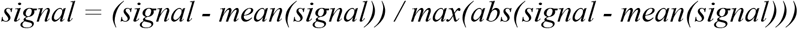

### Feature extraction and selection

We first ran a feature extraction algorithm to reduce redundancy of information and computational efforts in classifying the calls and to maximize the generalization ability of the classifier (Tajiri, Yabuwaki, Kitamura, & Abe, 2010). A popular method is extraction of mel frequency cepstral coefficients (MFCCs) (Supplementary Figure 2), a procedure that adapts function parameters to the primate auditory system (Fedurek, Zuberbühler, & Dahl, 2016; Mielke & Zuberbühler, 2013). While a typical spectrogram linearly scales frequencies (i.e., each frequency bin is spaced an equal number of Hertz apart), the mel-frequency scale is a logarithmical spacing of frequencies. We divided the calls into segments of 25ms length and 10ms steps between two successive segments. We warped 26 spectral bands and returned 13 cepstra, which resulted in feature dimensions of 13 values each. We then took the mean and co-variances of each cepstra over the collection of feature segments, resulting in a 13-value vector and a 13 x 13-value matrix, respectively, and concatenated to 104-unit vectors ((Mandel & Ellis, 2005), p. 594-599) (Figure 3). We applied feature scaling to [0 to 1] and mean normalization.

Second, we performed a feature selection procedure, a crucial part in statistical learning: too many feature dimensions are not useful for producing reliable classification systems, whereas low sample numbers can lead to over-fitting to noisy feature dimensions. We therefore selected a subset of the original feature dimensions and evaluated classification performance based on sequentially selected feature sets until there was no improvement in performance. At this end, we subdivided the entire data set into a training (75%) and a test data set (25%) and applied a *t*-test on each feature dimension, comparing values of given feature dimension sorted by predefined class labels (e.g. grunts occurring in negative (1) vs. positive (2) contexts) and used *p*-values as a measure separability of the two classes. We plotted the *p*-values as an empirical cumulative distribution function (eCDF) to get an understanding of how well each feature separated the two classes and how many features contributed to a significant separation (5%-level). We ran this procedure 20 times for each comparison and plotted the results individually (grey lines) and the mean of all repetitions (black line) (Figure 2A). The classification routines were then independently run either on feature dimensions selected according to the discrimination power (decreasing order) (red lines in Figure 2B), as shown in the eCDF plots (Figure 2A). Such procedure is referred to as a simple filter approach on feature selection, where general characteristics of the extracted features are taken into consideration when selecting feature dimensions, without subjecting them to a classifier. We also applied a more extensive procedure of feature selection by sequentially selecting feature dimensions by adding (forward search) feature dimensions, referred to as sequential feature selection (blue lines in Figure 2B). As part of this method, the algorithm searched the best feature dimensions (predictors) according to their individual classification performance in the given subset of data. For each candidate feature subset (predictor), the algorithm performed a 10-fold cross-validation procedure with different training and test subsets. After computing the mean performance values for each candidate feature subset, the algorithm chooses the candidate feature subset with minimal misclassification. For both methods, we systematically varied the number of features used for classification (x-axis in Figure 2B). The selected features from a single run of the sequential search algorithm are illustrated in Figure 2C. Scales reflect the feature-scaled and normalized values, as a result of feature extraction, from which the grand means (i.e. for each feature dimensions across all data) were subtracted. This measure was used to visually highlight differences and was not used in further analyses.

### Classification

We used support vector machine (SVM) with a radial basis function (RBF) Kernel (Vert, Tsuda, & Schölkopf, 2004) for the classification of calls according to the class labels (negative, neutral and positive contexts). A classification procedure contains a training phase followed by a test phase. We therefore separated training samples and labelled them according to an attribute of interest (e.g. negative (1) vs. positive (2) contexts). The algorithm then created a model that optimally separates the two classes. In the test phase, samples without attribute labels were fed into the model to measure its generalization performance. We used the SVM implementation from LIBSVM toolbox (Chang & Lin, 2011). To evaluate how the classification results generalize to a novel and independent data set, we 10-fold cross-validated the classification process and optimized the parameters C and gamma (Fedurek et al., 2016), with the C taking values in a range of (2^-1^, 2^3^) and gamma in a range of [2^-4^, 2^1^]. In addition, to ensure that no single individuals contributed solely to the classification outcome, we ran a leave-one-out algorithm, where the procedure described above was re-run six times, excluding one of the individuals in each run.

### Feature evaluation

To evaluate whether certain feature dimensions are particularly critical for the classification of grunts, we assessed whether feature dimensions have been repeatedly used by the classifier overall in the classification of grunts. We therefore considered the three types of comparisons, positive vs neutral, positive vs negative and neutral vs negative grunts, as well as the two feature evaluation algorithms (simple feature selection and sequential feature selection). Each comparison, as described above, was ten-fold cross-validated. We then calculated the empirical distribution of the ten features with best classification power, as determined by the feature selection algorithms (see above). Also, we determined a random distribution of “best features” for each comparison by randomly selecting 10 out of 104 features. The frequency distribution across all comparisons were determined and 95% confidence intervals were calculated by running the procedure 1,000 times. We then traced back the significant feature dimensions to the underlying frequency bands in Hertz.

## ACKNOWLEDGMENTS

We thank UWA and UNCST for permission to conduct the study, Geoffrey Muhanguzi, Caroline Asiimwe and Sam Adue for their support in the field, Derry Taylor for critical comments on previous versions of the manuscript.

We are grateful to the Royal Zoological Society of Scotland for providing core funding to the Budongo Conservation Field Station.

The research was supported by a Fyssen Fellowship and British Academy Newton International Fellowship awarded to GD, funding from the European Union’s Seventh Framework Programme for research, technological development and demonstration (grant agreement no 283871), and the Swiss National Science Foundation (PZ00P3_154741) awarded to CDD.

## SUPPLEMENTARY INFORMATION

**Supplementary Figure 1.**
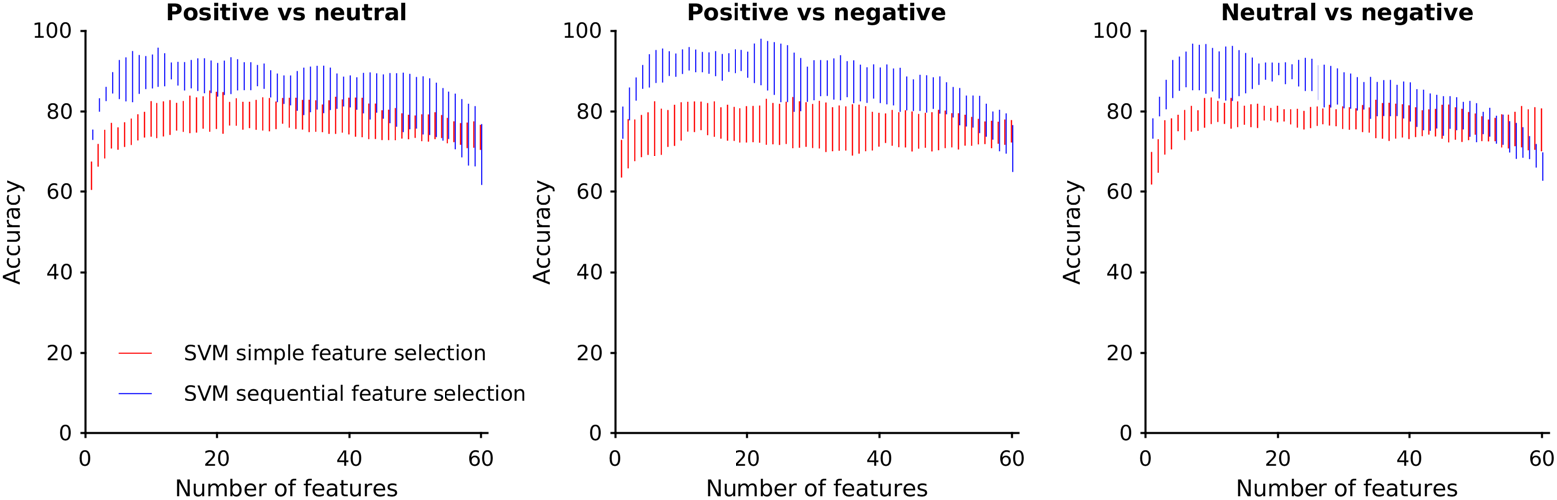
Leave-one-out method to account for subject effects. The accuracies of the three comparisons of grunt types are shown as function of number of features. These graphs illustrate the variability of accuracy caused by leaving out one of the 6 individuals per each separate classification procedure. The vertical bars indicate the minimum and maximum scores.

**Supplementary Figure 2.**
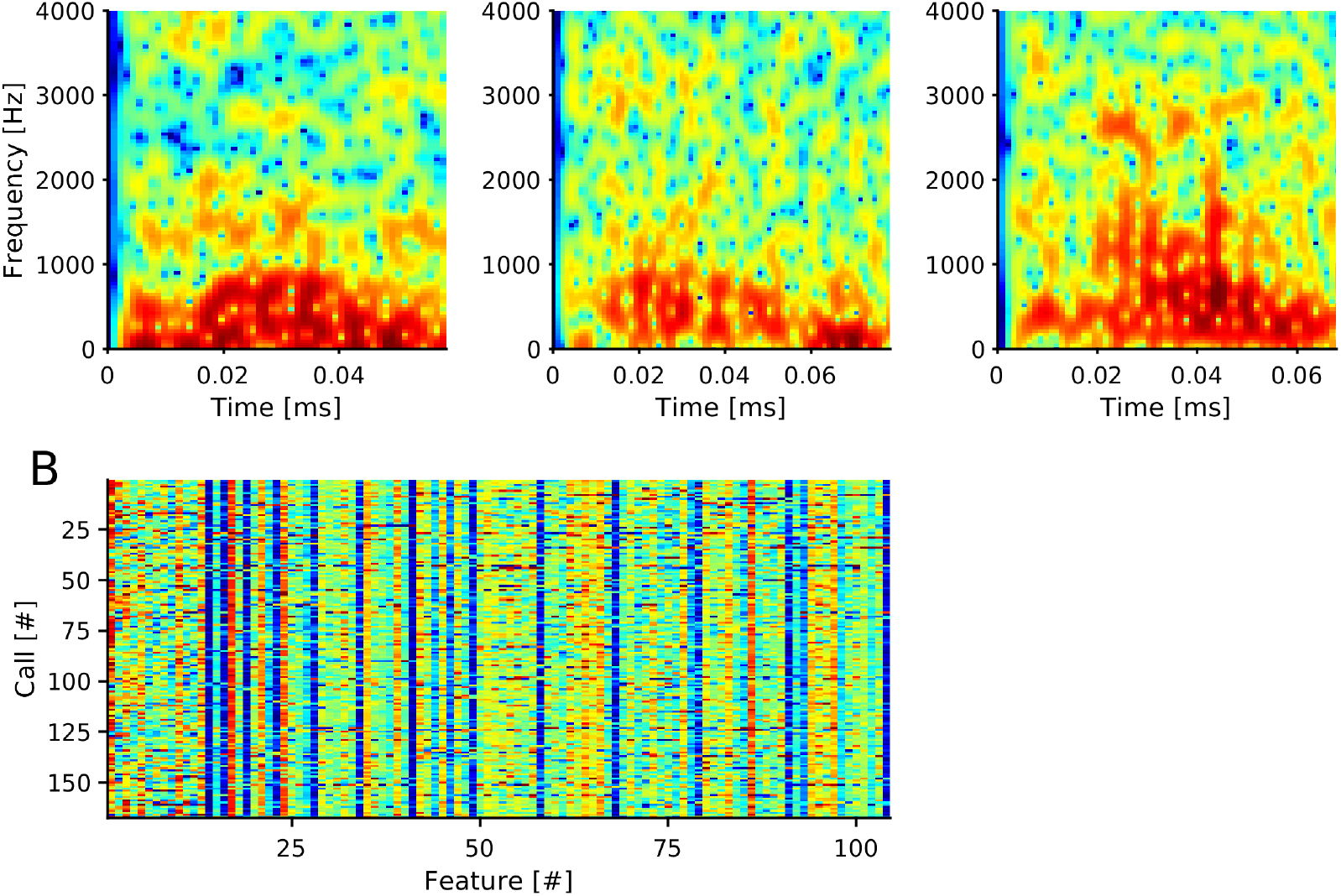
MFCCs extracted from example calls and extracted feature matrix. A. Time-frequency spectra of three arbitrarily chosen calls. B. From each call 26 spectral bands and 13 cepstra were extracted. Feature vectors containing the means and covariances of cepstra are shown for each call. Means are shown as features 1 to 13 on the x-axis, followed by covariances (91 values).

## REFERENCES

Arias, P., Belin, P., & Aucouturier, J.-J. (2018). Auditory smiles trigger unconscious facial imitation. Current Biology, 28(14), R782–R783. https://doi.org/10.1016/j.cub.2018.05.084

Aucouturier, J.-J., Johansson, P., Hall, L., Segnini, R., Mercadié, L., & Watanabe, K. (2016). Covert digital manipulation of vocal emotion alter speakers’ emotional states in a congruent direction. Proceedings of the National Academy of Sciences, 113(4), 948–953. https://doi.org/10.1073/pnas.1506552113

Banse, R., & Scherer, K. R. (1996). Acoustic profiles in vocal emotion expression. Journal of Personality and Social Psychology, 70(3), 614–636. https://doi.org/10.1037/0022-3514.70.3.614

Boë, L.-J., Berthommier, F., Legou, T., Captier, G., Kemp, C., Sawallis, T. R., … Fagot, J. (2017). Evidence of a Vocalic Proto-System in the Baboon (Papio papio) Suggests Pre-Hominin Speech Precursors. PLOS ONE, 12(1), e0169321. https://doi.org/10.1371/journal.pone.0169321

Boysson-Bardies, B. de. (2001). How Language Comes to Children: From Birth to Two Years. Cambridge, MA: MIT Press.

Briefer, E. F. (2012). Vocal expression of emotions in mammals: mechanisms of production and evidence. Journal of Zoology, 288(1), 1–20. https://doi.org/10.1111/j.1469-7998.2012.00920.x

Burkart, J. M., Hrdy, S. B., & Van Schaik, C. P. (2009). Cooperative breeding and human cognitive evolution. Evolutionary Anthropology, 18(5), 175–186.

Chang, C.-C., & Lin, C.-J. (2011). LIBSVM: a library for support vector machines. ACM Transactions on Intelligent Systems and Technology (TIST), 2(3), 27. https://doi.org/10.1145/1961189.1961199

Cheney, D. L., & Seyfarth, R. M. (1982). How vervet monkeys perceive their grunts: Field playback experiments. Animal Behaviour, 30(3), 739–751. https://doi.org/10.1016/S0003-3472(82)80146-2

Cheney, D. L., & Seyfarth, R. M. (2018). Flexible usage and social function in primate vocalizations. Proceedings of the National Academy of Sciences, 115(9), 1974–1979. https://doi.org/10.1073/pnas.1717572115

Clay, Z., Archbold, J., & Zuberbühler, K. (2015). Functional flexibility in wild bonobo vocal behaviour. PeerJ, 3, e1124. https://doi.org/10.7717/peerj.1124

Crockford, C. (in press). Why Does the Chimpanzee Vocal Repertoire Remain Poorly Understood? And What Can Be Done About It. In The Tai Chimpanzees: 40 years of Research. Eds: Boesch C. and Wittig R. Cambridge University Press.

Crockford, C., & Boesch, C. (2005). Call combinations in wild chimpanzees. Behaviour, 142(4), 397–421. https://doi.org/10.1163/1568539054012047

de Boysson-Bardies, B. (1993). Ontogeny of Language-Specific Syllabic Productions. In B. de Boysson-Bardies, S. de Schonen, P. Jusczyk, P. McNeilage, & J. Morton (Eds.), Developmental Neurocognition: Speech and Face Processing in the First Year of Life (pp. 353–363). https://doi.org/10.1007/978-94-015-8234-6_29

de Boysson-Bardies, B., & Vihman, M. M. (1991). Adaptation to Language: Evidence from Babbling and First Words in Four Languages. Language, 67(2), 297–319. https://doi.org/10.2307/415108

Dezecache, G., Zuberbühler, K., Davila-Ross, M., & Dahl, C. D. (2019). Machine learning reveals adaptive maternal responses to infant distress calls in wild chimpanzees. BioRxiv, 835827. https://doi.org/10.1101/835827

Elbers, L., & Ton, J. (1985). Play pen monologues: the interplay of words and babbles in the first words period. Journal of Child Language, 12(3), 551–565. https://doi.org/10.1017/S0305000900006644

Fedurek, P., Zuberbühler, K., & Dahl, C. D. (2016). Sequential information in a great ape utterance. Scientific Reports, 6, 38226. https://doi.org/10.1038/srep38226

Fischer, J. (2016). Playback Experiments. The International Encyclopedia of Primatology, 1–2. https://doi.org/10.1002/9781119179313.wbprim0140

Fischer, J., Noser, R., & Hammerschmidt, K. (2013). Bioacoustic field research: a primer to acoustic analyses and playback experiments with primates. American Journal of Primatology, 75(7), 643–663. https://doi.org/10.1002/ajp.22153

Fitch, W. T. (2018). The Biology and Evolution of Speech: A Comparative Analysis. Annual Review of Linguistics, 4(1), 255–279. https://doi.org/10.1146/annurev-linguistics-011817-045748

Fitch, W. T., Boer, B. de, Mathur, N., & Ghazanfar, A. A. (2016). Monkey vocal tracts are speech-ready. Science Advances, 2(12), e1600723. https://doi.org/10.1126/sciadv.1600723

Fröhlich, M., & Hobaiter, C. (2018). The development of gestural communication in great apes. Behavioral Ecology and Sociobiology, 72(12), 194. https://doi.org/10.1007/s00265-018-2619-y

Fröhlich, M., Wittig, R. M., & Pika, S. (2019). The ontogeny of intentional communication in chimpanzees in the wild. Developmental Science, e12716. doi: 10.1111/desc.12716

Ghazanfar, A. A., Liao, D. A., & Takahashi, D. Y. (2019). Volition and learning in primate vocal behaviour. Animal Behaviour, 151, 239–247. https://doi.org/10.1016/j.anbehav.2019.01.021

Ghazanfar, A. A., & Takahashi, D. Y. (2014). The evolution of speech: vision, rhythm, cooperation. Trends in Cognitive Sciences, 18(10), 543–553. https://doi.org/10.1016/j.tics.2014.06.004

Goodall, J. (1986). The chimpanzees of Gombe: Patterns of behavior. Cambridge, MA: Harvard University Press.

Goupil, L., Johansson, P., Hall, L., & Aucouturier, J.-J. (2019). Influence of Vocal Feedback on Emotions Provides Causal Evidence for the Self-Perception Theory. https://doi.org/10.1101/510867

Hayes, K. J., & Hayes, C. (1951). The Intellectual Development of a Home-Raised Chimpanzee. Proceedings of the American Philosophical Society, 95(2), 105–109.

Hrdy, S. (2007). Evolutionary context of human development: the cooperative breeding model. In Family Relationships: An Evolutionary Perspective. Oxford: Oxford University Press.

Hrdy, S. (2009). Mothers and others. Cambridge, MA: Harvard University Press.

Hsu, M. J., Chen, L.-M., & Agoramoorthy, G. (2005). The vocal repertoire of Formosan macaques, Macaca cyclopis: acoustic structure and behavioral context. Zoological Studies, 44(2), 275.

Jhang, Y., & Oller, D. K. (2017). Emergence of Functional Flexibility in Infant Vocalizations of the First 3 Months. Frontiers in Psychology, 8. https://doi.org/10.3389/fpsyg.2017.00300

Kersken, V., Zuberbühler, K., & Gomez, J.-C. (2017). Listeners can extract meaning from non-linguistic infant vocalisations cross-culturally. Scientific Reports, 7, srep41016. https://doi.org/10.1038/srep41016

Kojima, S. (2003). A search for the origins of human speech: Auditory and vocal functions of the chimpanzee. Kyoto: Kyoto University Academic Press.

Kramer, K. L. (2010). Cooperative Breeding and its Significance to the Demographic Success of Humans. Annual Review of Anthropology, 39(1), 417–436. https://doi.org/10.1146/annurev.anthro.012809.105054

Lameira, A. R., & Call, J. (2018). Time-space-displaced responses in the orangutan vocal system. Science Advances, 4(11), eaau3401. https://doi.org/10.1126/sciadv.aau3401

Lameira, A. R., & Shumaker, R. W. (2019). Orangutans show active voicing through a membranophone. Scientific Reports., 9, 12289. https://doi.org/10.1038/s41598-019-48760-7

Laporte, M. N. C., & Zuberbühler, K. (2011). The development of a greeting signal in wild chimpanzees. Developmental Science, 14(5), 1220–1234. https://doi.org/10.1111/j.1467-7687.2011.01069.x

Levréro, F., & Mathevon, N. (2013). Vocal Signature in Wild Infant Chimpanzees. American Journal of Primatology, 75(4), 324–332. https://doi.org/10.1002/ajp.22108

Lieberman, P. (2017). Comment on “Monkey vocal tracts are speech-ready.” Science Advances, 3(7), e1700442. https://doi.org/10.1126/sciadv.1700442

Locke, J. L. (2006). Parental selection of vocal behavior. Human Nature, 17(2), 155–168. https://doi.org/10.1007/s12110-006-1015-x

Mandel, M. I., & Ellis, D. P. (2005). Song-level features and support vector machines for music classification. Proceedings of the 6th International Conference on Music Information Retrieval (ISMIR), 594–599.

McCune, L., Vihman, M. M., Roug-Hellichius, L., Delery, D. B., & Gogate, L. (1996). Grunt communication in human infants (Homo sapiens). Journal of Comparative Psychology, 110(1), 27.

Mielke, A., & Zuberbühler, K. (2013). A method for automated individual, species and call type recognition in free-ranging animals. Animal Behaviour, 86(2), 475–482. https://doi.org/10.1016/j.anbehav.2013.04.017

Møller, A. P. (1988). False Alarm Calls as a Means of Resource Usurpation in the Great Tit Parus major. Ethology, 79(1), 25–30. https://doi.org/10.1111/j.1439-0310.1988.tb00697.x

Nathani, S., Ertmer, D. J., & Stark, R. E. (2006). Assessing vocal development in infants and toddlers. Clinical Linguistics & Phonetics, 20(5), 351–369. https://doi.org/10.1080/02699200500211451

Oller, D. K. (2000). The Emergence of the Speech Capacity. https://doi.org/10.4324/9781410602565

Oller, D. K., Buder, E. H., Ramsdell, H. L., Warlaumont, A. S., Chorna, L., & Bakeman, R. (2013). Functional flexibility of infant vocalization and the emergence of language. Proceedings of the National Academy of Sciences of the United States of America, 110(16), 6318–6323. https://doi.org/10.1073/pnas.1300337110

Oller, D. K., & Griebel, U. (2004). Contextual freedom in human infant vocalization and the evolution of language. In Evolution of communicative flexibility: Complexity, creativity and adaptability in human and animal communication (p. 135). Cambridge, MA: MIT Press.

Oller, D. K., Griebel, U., Iyer, S. N., Jhang, Y., Warlaumont, A. S., Dale, R., & Call, J. (2019). Language Origins Viewed in Spontaneous and Interactive Vocal Rates of Human and Bonobo Infants. Frontiers in Psychology, 10. https://doi.org/10.3389/fpsyg.2019.00729

Oller, D. K., Griebel, U., & Warlaumont, A. S. (2016). Vocal Development as a Guide to Modeling the Evolution of Language. Topics in Cognitive Science, 8(2), 382–392. https://doi.org/10.1111/tops.12198

Oller, D. K., Wieman, L. A., Doyle, W. J., & Ross, C. (1976). Infant babbling and speech. Journal of Child Language, 3(1), 1–11. https://doi.org/10.1017/S0305000900001276

Owren, M. J., & Casale, T. M. (1994). Variations in fundamental frequency peak position in Japanese macaque (Macaca fuscata) coo calls. Journal of Comparative Psychology, 108(3), 291. https://doi.org/10.1037/0735-7036.108.3.291

Plooij, F. X. (1984). The behavioral development of free-living chimpanzee babies and infants. Norwood, NJ: Ablex.

Pollick, A. S., & Waal, F. B. M. de. (2007). Ape gestures and language evolution. Proceedings of the National Academy of Sciences, 104(19), 8184–8189. https://doi.org/10.1073/pnas.0702624104

Ponsot, E., Burred, J. J., Belin, P., & Aucouturier, J.-J. (2018). Cracking the social code of speech prosody using reverse correlation. Proceedings of the National Academy of Sciences, 115(15), 3972–3977. https://doi.org/10.1073/pnas.1716090115

R Core Team. (2018). R: A language and environment for statistical computing. Retrieved from https://www.R-project.org/

Range, F., & Fischer, J. (2004). Vocal Repertoire of Sooty Mangabeys (Cercocebus torquatus atys) in the Taï National Park. Ethology, 110(4), 301–321. https://doi.org/10.1111/j.1439-0310.2004.00973.x

Rendall, D., Seyfarth, R. M., Cheney, D. L., & Owren, M. J. (1999). The meaning and function of grunt variants in baboons. Animal Behaviour, 57(3), 583–592. https://doi.org/10.1006/anbe.1998.1031

Rendall, D., & Owren, M. J. (2002). Animal vocal communication: Say what? In The cognitive animal: Empirical and theoretical perspectives on animal cognition (pp. 307–313). Cambridge, MA, US: MIT Press.

Reynolds, V. (2005). The chimpanzees of the Budongo Forest: ecology, behaviour, and conservation. Retrieved from http://books.google.fr/books?hl=fr&lr=&id=C6hzM5lQJ6YC&oi=fnd&pg=PR11&dq=budongo+reynolds&ots=OOfJtMfycP&sig=X1c6kEzGs8ZlzhXdNH3aK3foPrA

RStudio Team. (2015). RStudio: integrated development for R. RStudio, Inc., Boston, MA URL Http://Www.Rstudio.Com, 42.

Salmi, R., Hammerschmidt, K., & Doran-Sheehy, D. M. (2013). Western Gorilla Vocal Repertoire and Contextual Use of Vocalizations. Ethology, 119(10), 831–847. https://doi.org/10.1111/eth.12122

Schaik, C. P. van, & Burkart, J. M. (2010). Mind the Gap: Cooperative Breeding and the Evolution of Our Unique Features. In Mind the Gap (pp. 477–496). https://doi.org/10.1007/978-3-642-02725-3_22

Seyfarth, R. M., & Cheney, D. L. (1984). The acoustic features of vervet monkey grunts. The Journal of the Acoustical Society of America, 75(5), 1623–1628. https://doi.org/10.1121/1.390872

Slocombe, K. E., & Zuberbühler, K. (2010). Vocal communication in chimpanzees. The Mind of the Chimpanzee: Ecological and Experimental Perspectives. University of Chicago Press, Chicago, 192–207.

Slocombe, K. E., & Zuberbühler, K. (2005). Functionally Referential Communication in a Chimpanzee. Current Biology, 15(19), 1779–1784. https://doi.org/10.1016/j.cub.2005.08.068

Tajiri, Y., Yabuwaki, R., Kitamura, T., & Abe, S. (2010). Feature Extraction Using Support Vector Machines. In K. W. Wong, B. S. U. Mendis, & A. Bouzerdoum (Eds.), Neural Information Processing. Models and Applications (pp. 108–115). https://doi.org/10.1007/978-3-642-17534-3_14

Takahashi, D. Y., Fenley, A. R., Teramoto, Y., Narayanan, D. Z., Borjon, J. I., Holmes, P., & Ghazanfar, A. A. (2015). The developmental dynamics of marmoset monkey vocal production. Science, 349(6249), 734–738. https://doi.org/10.1126/science.aab1058

Vert, J.-P., Tsuda, K., & Schölkopf, B. (2004). A primer on kernel methods. In Kernel methods in computational biology (Vol. 47, pp. 35–70). Cambridge, MA: MIT Press.

Waal, F. B. M. de, & Pollick, A. S. (2011). Gesture as the most flexible modality of primate communication. The Oxford Handbook of Language Evolution. https://doi.org/10.1093/oxfordhb/9780199541119.013.0006

Watson, S. K., Townsend, S. W., Schel, A. M., Wilke, C., Wallace, E. K., Cheng, L.,. Slocombe, K. E. (2015). Vocal Learning in the Functionally Referential Food Grunts of Chimpanzees. Current Biology, 25(4), 495–499. https://doi.org/10.1016/j.cub.2014.12.032

Wheeler, B. C. (2009). Monkeys crying wolf? Tufted capuchin monkeys use anti-predator calls to usurp resources from conspecifics. Proceedings of the Royal Society B: Biological Sciences, 276(1669), 3013–3018. https://doi.org/10.1098/rspb.2009.0544

Williams, C. E., & Stevens, K. N. (1972). Emotions and Speech: Some Acoustical Correlates. The Journal of the Acoustical Society of America, 52(4B), 1238–1250. https://doi.org/10.1121/1.1913238

Yoo, H., Bowman, D. A., & Oller, D. K. (2018). The Origin of Protoconversation: An Examination of Caregiver Responses to Cry and Speech-Like Vocalizations. Frontiers in Psychology, 9. https://doi.org/10.3389/fpsyg.2018.01510

Zuberbühler, K. (2012). Cooperative breeding and the evolution of vocal flexibility. In The Oxford Handbook of Language Evolution. Oxford: Oxford University Press.

Zuberbühler, K. (2014). Experimental field studies with non-human primates. Current Opinion in Neurobiology, 28, 150–156. https://doi.org/10.1016/j.conb.2014.07.012

